# Functional consequences of genetic loci associated with intelligence in a meta-analysis of 87,740 individuals

**DOI:** 10.1101/170712

**Authors:** Jonathan R. I. Coleman, Julien Bryois, Héléna A. Gaspar, Philip R. Jansen, Jeanne Savage, Nathan Skene, Robert Plomin, Ana B. Muñoz-Manchado, Sten Linnarsson, Greg Crawford, Jens Hjerling-Leffler, Patrick F. Sullivan, Danielle Posthuma, Gerome Breen

## Abstract

Variance in IQ is associated with a wide range of health outcomes, and 1% of the population are affected by intellectual disability. Despite a century of research, the fundamental neural underpinnings of intelligence remain unclear. We integrate results from genome-wide association studies (GWAS) of intelligence with brain tissue and single cell gene expression data to identify tissues and cell types associated with intelligence. GWAS data for IQ (N = 78,308) were meta-analyzed with an extreme-trait cohort of 1,247 individuals with mean IQ ∼170 and 8,185 controls. Genes associated with intelligence implicate pyramidal neurons of the somatosensory cortex and CA1 region of the hippocampus, and midbrain embryonic GABAergic neurons. Tissue-specific analyses find the most significant enrichment for frontal cortex brain expressed genes. These results suggest specific neuronal cell types and genes may be involved in intelligence and provide new hypotheses for neuroscience experiments using model systems.

---

Genome-wide association studies (GWAS) have successfully identified statistical associations with a wide range of behavioral phenotypes and neuropsychiatric disorders^1–3^. Increasing sample sizes has begun to yield findings for intelligence^4–6^. The largest and most recent study of intelligence reported 18 loci significantly associated with intelligence^4^. Significant genetic correlations were observed between intelligence and a variety of behavioral (educational attainment, smoking behaviors), anthropometric (cranial morphology, height, body composition), and psychiatric phenotypes (schizophrenia, autism, depressive symptoms), mirroring epidemiological evidence for correlations between intelligence and a broad range of health-related outcomes^4,7^.

Considered alone, not all associations identified by GWAS precisely localize biological mechanisms amenable to subsequent experimentation. For instance, the most associated variant in a significant locus may not be the causal variant^8,9^, there may be multiple causal variants in a locus^10^, loci may act through altering the expression of distant genes^11^, and the associations identified by GWAS of complex traits are often spread across the genome^12–14^. To extract meaningful biological inferences from GWAS results, it is necessary to integrate data from other sources, such as studies of gene expression^15^. Results from gene-wise analyses in Sniekers et al. (2017) identified expression predominant in the brain for 14 of the 44 genes with significant association, although some transcription was inferred for most genes across most tissues^4^. Integration of genomic results with data on biological pathways suggested a prominent role for nervous system development in intelligence.

*In silico* functional annotation of GWAS results is dependent on high-quality biological reference data. Recently, data from the Karolinska Institutet (KI) mouse superset of single-cell RNA sequencing (scRNAseq) of ∼10,000 single cells from multiple brain regions was used to map schizophrenia GWAS results to brain cell types^16^. Genes that previously showed association with schizophrenia were expressed with higher specificity in pyramidal cells, medium spiny neurons, and interneurons than in 20 other brain cell types. This demonstrates the potential of cell-type specific annotation to enable the construction of new functional hypotheses for complex traits.

We sought to develop a better understanding of the neurobiological underpinnings of intelligence through combining GWAS results with a number of data sources concerned with tissue and cell-type specific gene expression. To this end, we meta-analyzed the most recent GWAS of intelligence^4^ with an extreme-trait GWAS that compared individuals of very high intelligence to a group from the general population (HiQ)^17^. We then analyzed the enrichment of associations with intelligence in single cell expression data from the KI mouse superset^16^, and in brain genomic and transcriptomic data from the GTEx project^18^. Finally, we combined genomic and tissue-specific expression data with information from biological, disease-relevant and drug-target pathway databases to further assess the potential impact of biological mechanisms explaining variance in intelligence.

## Results

### Meta-analysis

The genetic correlation between Sniekers et al. (2017) and HiQ was 0.92 (SE: 0.07) and was sufficiently high to justify meta-analysis. 25 loci met genome-wide significance with intelligence (i.e., p < 5 × 10^-8^; Table 1, Figure 1a). Of these, 13 were genome-wide significant in Sniekers et al. (2017) and 12 were novel (Table 1; Supplementary Table 1). The single locus previously identified in the HiQ sample was not genome-wide significant in this analysis (p = 0.00595;^17^).

**Table 1:**
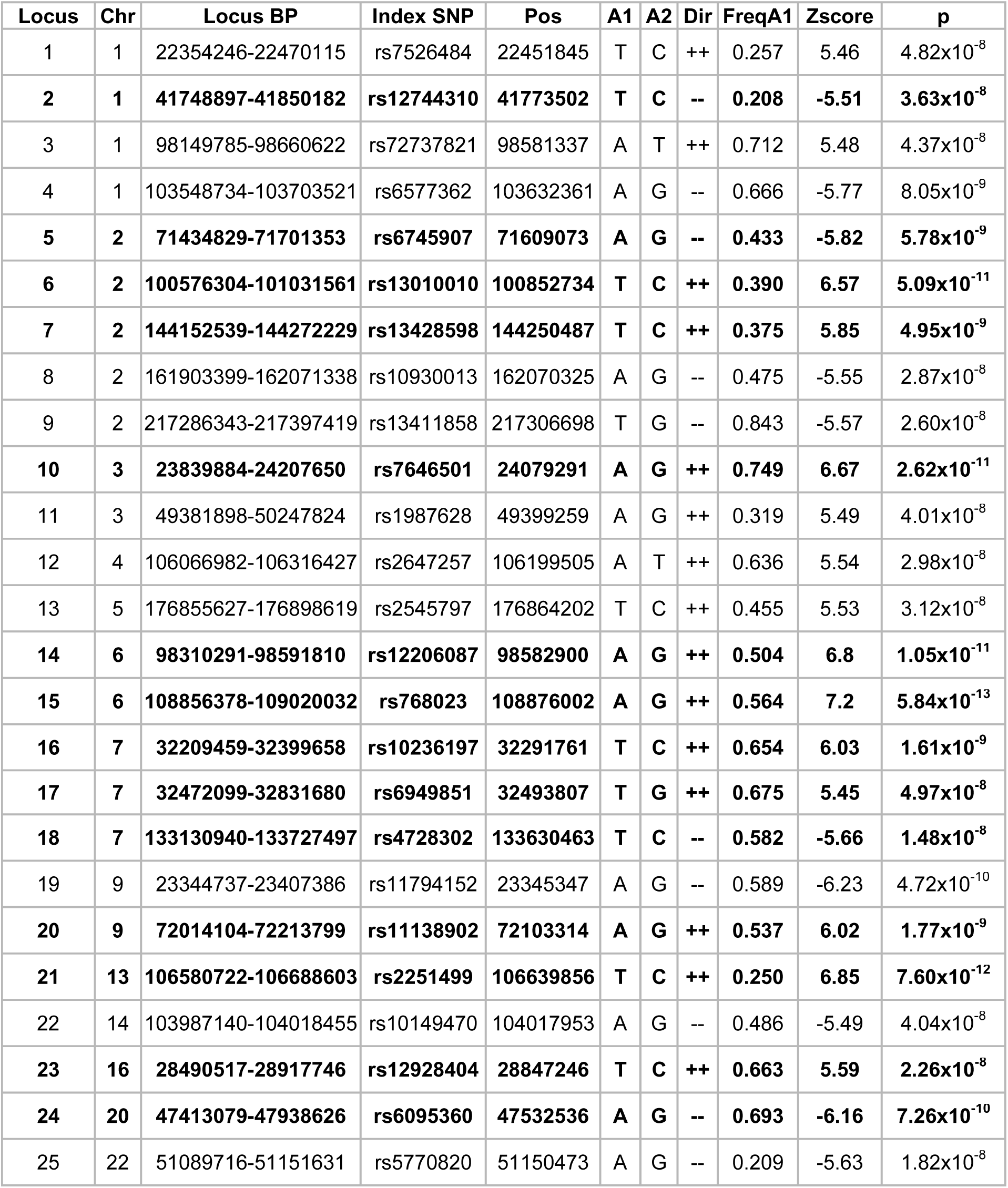
Genome-wide significant variants from single-variant association analyses. Loci significant in Sniekers et al. (2017) are in bold. FreqA1 = A1 frequency in non-Finnish samples from the 1000 Genomes project. Dir = direction of effect in Sniekers and in TIP.

| Locus | Chr | Locus BP | Index SNP | Pos | A1 | A2 | Dir | FreqA1 | Zscore | p |
| --- | --- | --- | --- | --- | --- | --- | --- | --- | --- | --- |
| 1 | 1 | 22354246-22470115 | rs7526484 | 22451845 | T | C | ++ | 0.257 | 5.46 | 4.82x10<sup>-8</sup> |
| **2** | **1** | **41748897-41850182** | **rs12744310** | **41773502** | **T** | **C** | **--** | **0.208** | **-5.51** | **3.63x10<sup>-8</sup>** |
| 3 | 1 | 98149785-98660622 | rs72737821 | 98581337 | A | T | ++ | 0.712 | 5.48 | 4.37x10<sup>-8</sup> |
| 4 | 1 | 103548734-103703521 | rs6577362 | 103632361 | A | G | -- | 0.666 | -5.77 | 8.05x10<sup>-9</sup> |
| **5** | **2** | **71434829-71701353** | **rs6745907** | **71609073** | **A** | **G** | **--** | **0.433** | **-5.82** | **5.78x10<sup>-9</sup>** |
| **6** | **2** | **100576304-101031561** | **rs13010010** | **100852734** | **T** | **C** | **++** | **0.390** | **6.57** | **5.09x10<sup>-11</sup>** |
| **7** | **2** | **144152539-144272229** | **rs13428598** | **144250487** | **T** | **C** | **++** | **0.375** | **5.85** | **4.95x10<sup>-9</sup>** |
| 8 | 2 | 161903399-162071338 | rs10930013 | 162070325 | A | G | -- | 0.475 | -5.55 | 2.87x10<sup>-8</sup> |
| 9 | 2 | 217286343-217397419 | rs13411858 | 217306698 | T | G | -- | 0.843 | -5.57 | 2.60x10<sup>-8</sup> |
| **10** | **3** | **23839884-24207650** | **rs7646501** | **24079291** | **A** | **G** | **++** | **0.749** | **6.67** | **2.62x10<sup>-11</sup>** |
| 11 | 3 | 49381898-50247824 | rs1987628 | 49399259 | A | G | ++ | 0.319 | 5.49 | 4.01x10<sup>-8</sup> |
| 12 | 4 | 106066982-106316427 | rs2647257 | 106199505 | A | T | ++ | 0.636 | 5.54 | 2.98x10<sup>-8</sup> |
| 13 | 5 | 176855627-176898619 | rs2545797 | 176864202 | T | C | ++ | 0.455 | 5.53 | 3.12x10<sup>-8</sup> |
| **14** | **6** | **98310291-98591810** | **rs12206087** | **98582900** | **A** | **G** | **++** | **0.504** | **6.8** | **1.05x10<sup>-11</sup>** |
| **15** | **6** | **108856378-109020032** | **rs768023** | **108876002** | **A** | **G** | **++** | **0.564** | **7.2** | **5.84x10<sup>-13</sup>** |
| **16** | **7** | **32209459-32399658** | **rs10236197** | **32291761** | **T** | **C** | **++** | **0.654** | **6.03** | **1.61x10<sup>-9</sup>** |
| **17** | **7** | **32472099-32831680** | **rs6949851** | **32493807** | **T** | **G** | **++** | **0.675** | **5.45** | **4.97x10<sup>-8</sup>** |
| **18** | **7** | **133130940-133727497** | **rs4728302** | **133630463** | **T** | **C** | **--** | **0.582** | **-5.66** | **1.48x10<sup>-8</sup>** |
| 19 | 9 | 23344737-23407386 | rs11794152 | 23345347 | A | G | -- | 0.589 | -6.23 | 4.72x10<sup>-10</sup> |
| **20** | **9** | **72014104-72213799** | **rs11138902** | **72103314** | **A** | **G** | **++** | **0.537** | **6.02** | **1.77x10<sup>-9</sup>** |
| **21** | **13** | **106580722-106688603** | **rs2251499** | **106639856** | **T** | **C** | **++** | **0.250** | **6.85** | **7.60x10<sup>-12</sup>** |
| 22 | 14 | 103987140-104018455 | rs10149470 | 104017953 | A | G | -- | 0.486 | -5.49 | 4.04x10<sup>-8</sup> |
| **23** | **16** | **28490517-28917746** | **rs12928404** | **28847246** | **T** | **C** | **++** | **0.663** | **5.59** | **2.26x10<sup>-8</sup>** |
| **24** | **20** | **47413079-47938626** | **rs6095360** | **47532536** | **A** | **G** | **--** | **0.693** | **-6.16** | **7.26x10<sup>-10</sup>** |
| 25 | 22 | 51089716-51151631 | rs5770820 | 51150473 | A | G | -- | 0.209 | -5.63 | 1.82x10<sup>-8</sup> |

**Figure 1:**
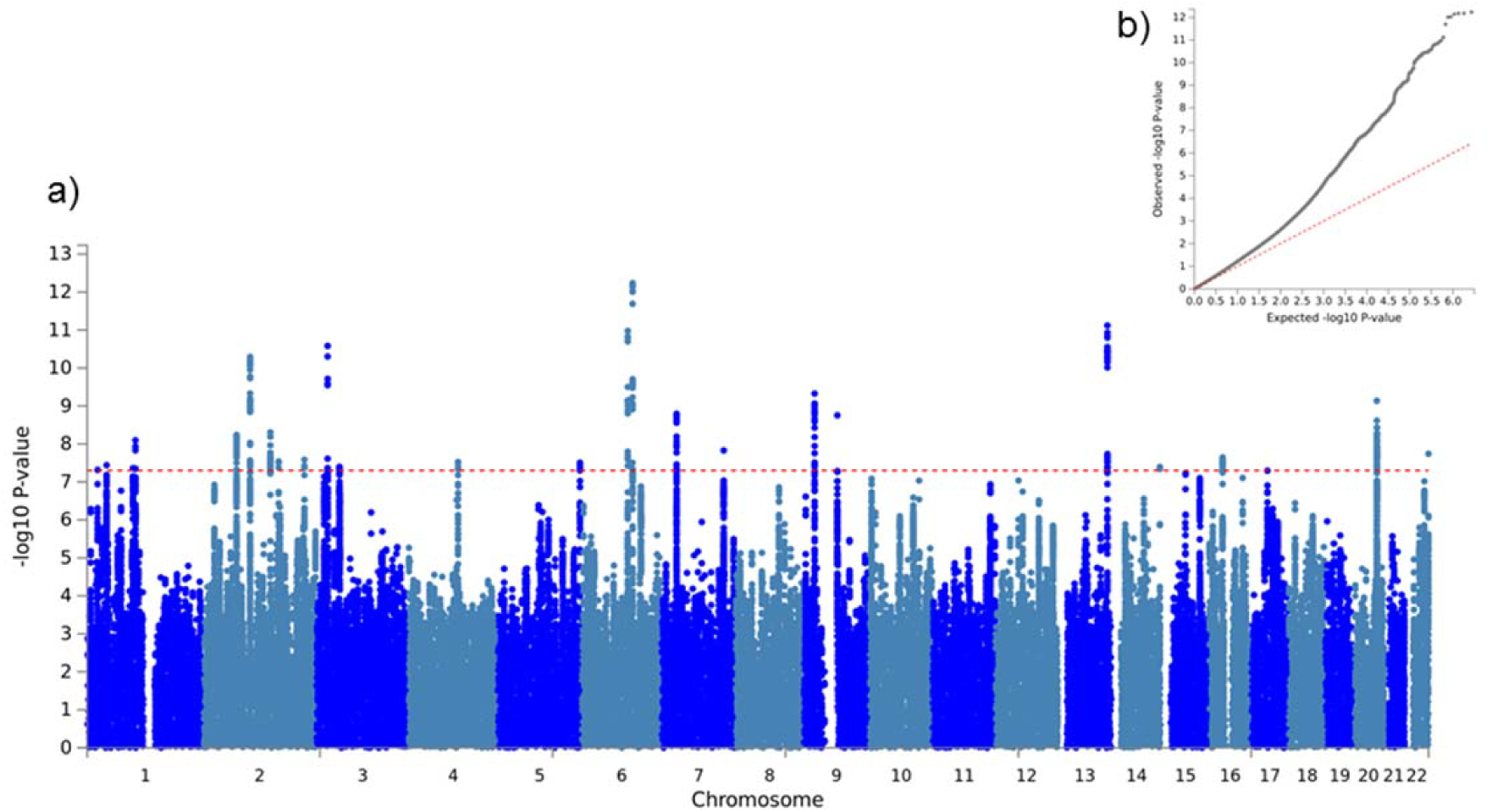
a) Manhattan plot and b) QQ plot of meta-analysis results

Assessment of genome-wide inflation yielded a median *λ*_GC_ of 1.24. The LD score regression intercept from was 1.004 (SE: 0.01), suggesting that this inflation is caused by polygenicity rather than confounding (Figure 1b)^19^. Annotation of specific genomic loci to databases of interest suggested overlapping loci between intelligence and a variety of phenotypes (Supplementary Table 2). The largest overlap was observed with educational attainment (14/25 loci), but overlap at multiple loci was widespread, including with age at menarche, height, body mass index, autoimmune disease, and schizophrenia. Genes previously implicated in intellectual disability or developmental delay (ID/DD) were present within 9/25 loci.

### Heritability and partitioned heritability

LD Score regression yielded a SNP-heritability estimate of 0.221 (SE: 0.01) in line with the previously reported SNP-heritability in Sniekers et al (2017). Partitioning this heritability across 58 functional SNP annotations identifies conserved regions as significantly enriched contributors to the heritability of intelligence (proportion of heritability = 0.340, enrichment = 13.3 fold, p = 3.26 × 10^-8^), consistent with previous reports in a subset of our meta-analyzed cohorts^20^. Four extended annotations were also significantly enriched (p < 8.62 × 10^-4^; Figure 2), suggesting that genetic variation located in the vicinity of conserved regions, enhancers (specifically H3K4me1 elements) and open chromatin in brain dorsolateral prefrontal cortex^21^ and fetal cells) are enriched in heritability for intelligence.

**Figure 2:**
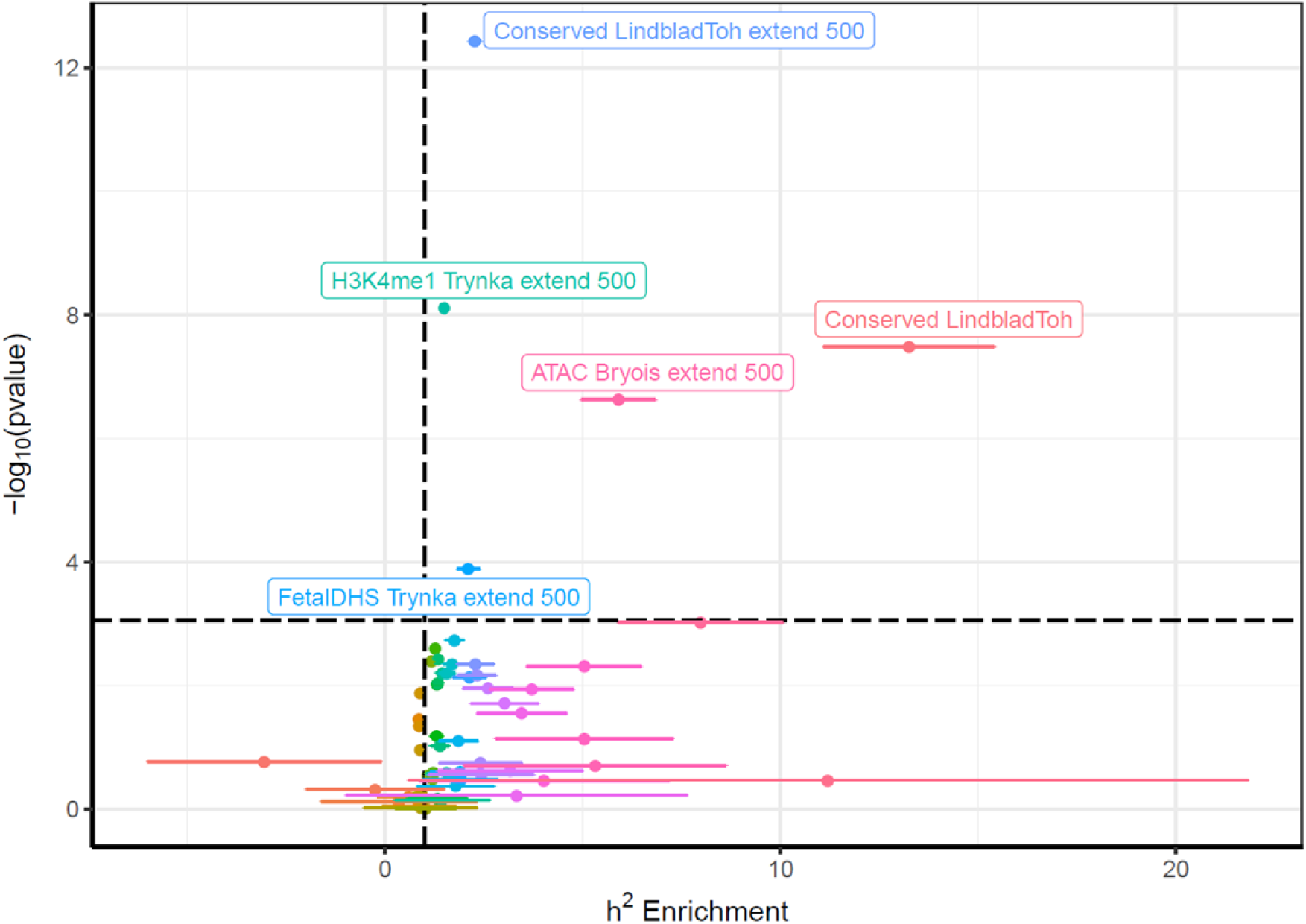
Heritability enrichment of genomic annotations. Horizontal line is p = 8.62×10^-4^, Bonferroni-corrected threshold for 58 annotations. Vertical line is enrichment = 1 (that is, no enrichment)

### Gene-wise analyses

93 genes at 41 loci were identified at genome-wide significance (p < 2.65 × 10^-6^, Bonferroni threshold for 18,839 genes; Table 2, Supplementary Figure 1). 28 of these genes (20 loci) were also identified in Sniekers et al. (2017). 11 of the 93 genes were previously implicated in intellectual disability or developmental delay, although this overlap does not significantly from what is expected under the null hypothesis of no enrichment (p = 0.073, hypergeometric test; Table 2).

**Table 2:**
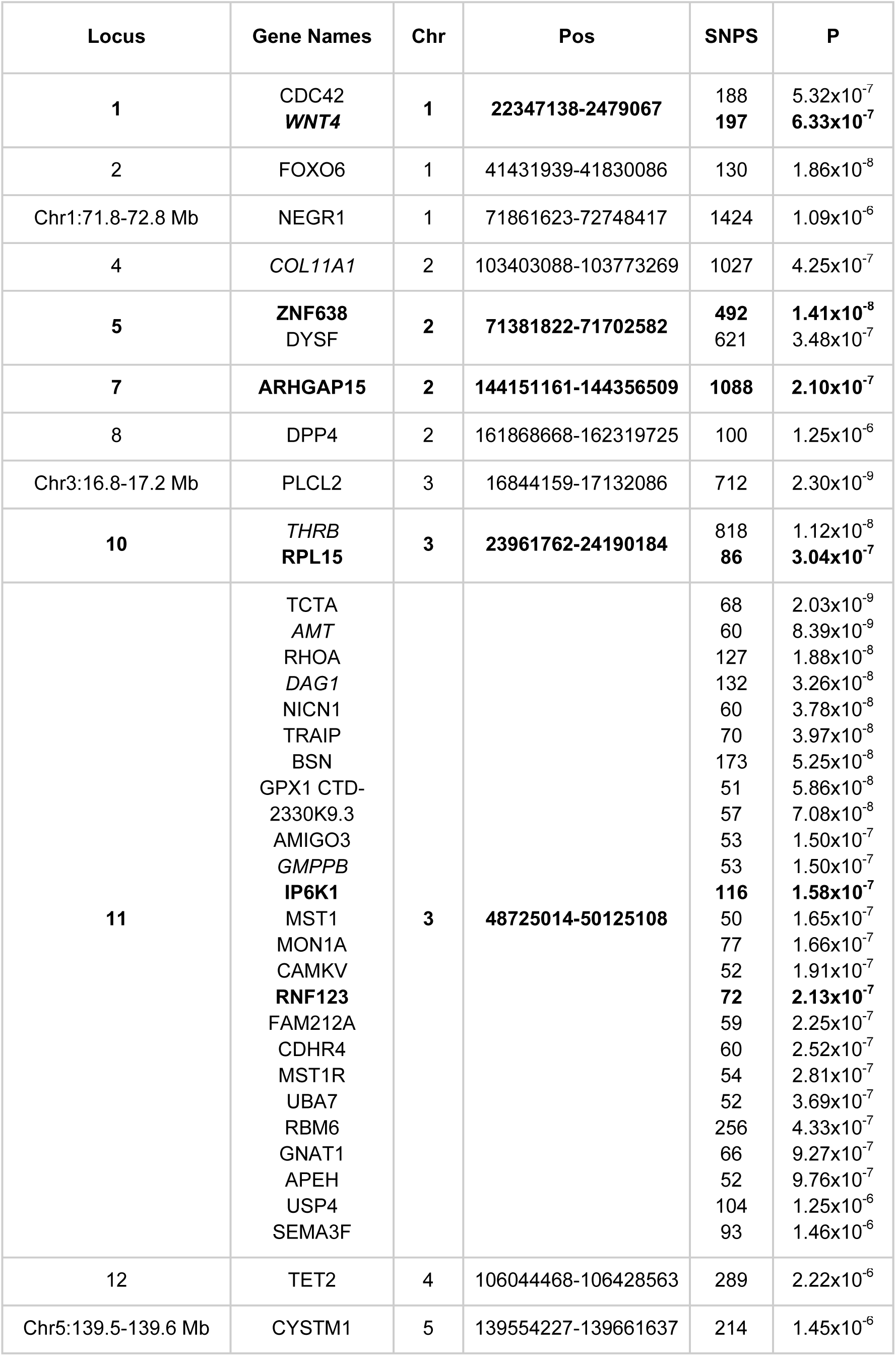
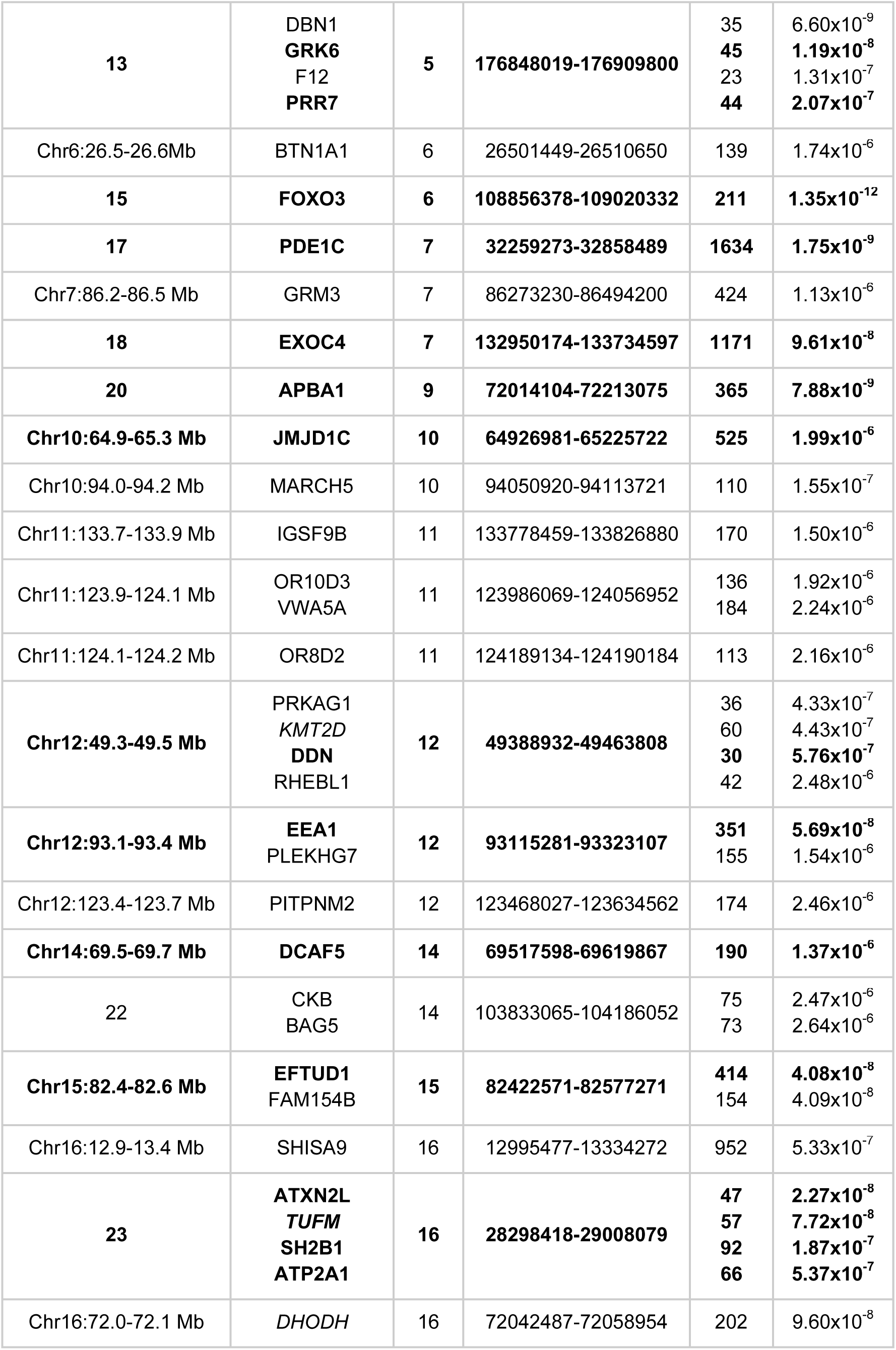
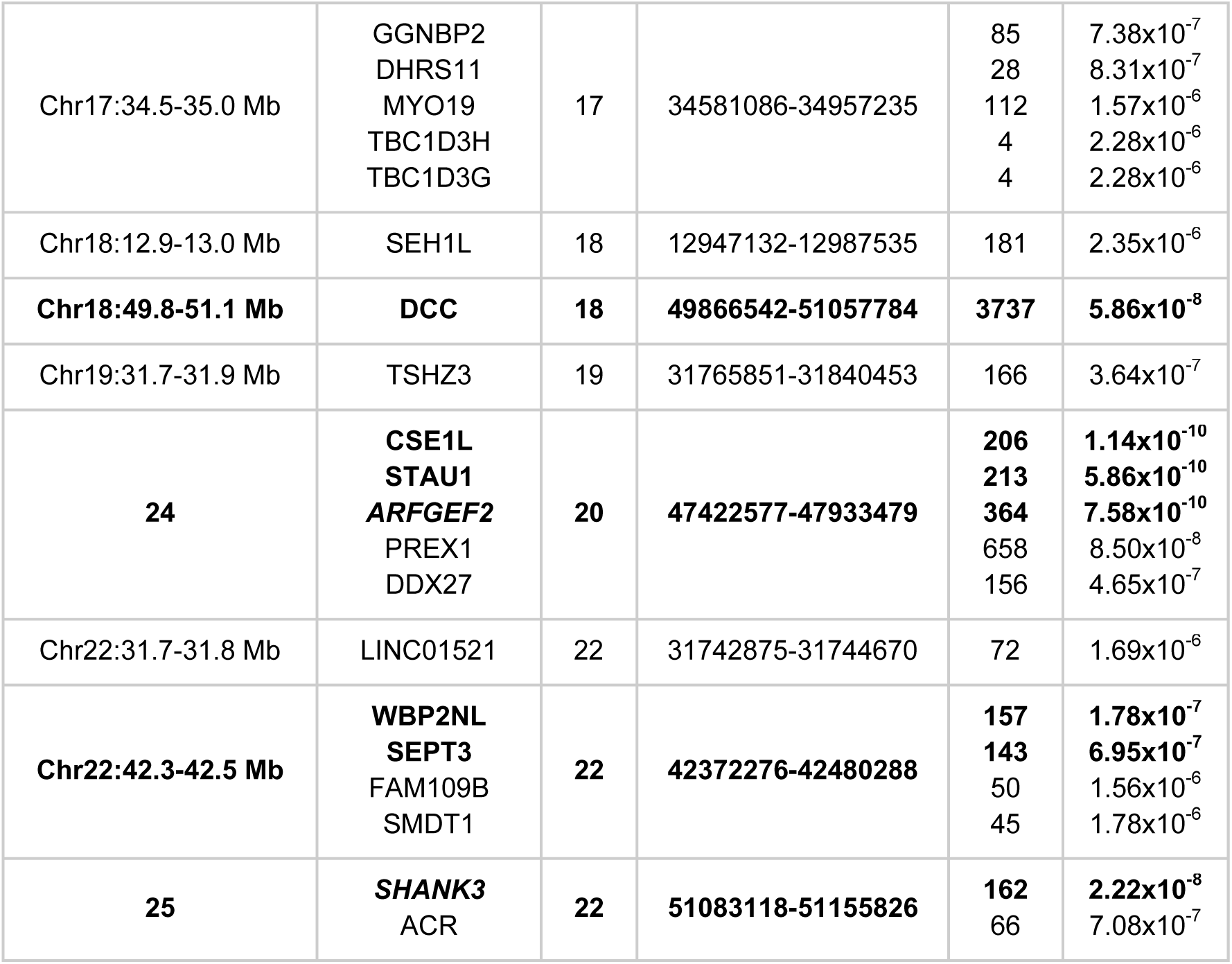
Genome-wide significant genes from gene-wise association analyses. Genetic locus is that identified in single-variant analyses, or the genomic region otherwise. P value from competitive test in MAGMA. Loci in bold were significantly associated in Sniekers et al. (2017). Genes in italics have previously been implicated in developmental delay or intellectual disability.

### Tissue- and cell-specific gene expression

Tissue-specific enrichment analysis identified an enrichment of genes with high brain-specific expression associated with intelligence (p_MAGMA_ = 4.43 × 10^-9^, p_LDSC_ = 4.23 × 10^-6^; Figure 3a, Supplementary Table 3a). Across 10 brain regions in GTEx^18^, stronger gene associations with intelligence were associated with increased specificity of gene expression to the frontal cortex (p_MAGMA_ = 0.00305, p_LDSC_ = 2.66 × 10^-4^; Figure 3b, Supplementary Table 3b).

**Figure 3:**
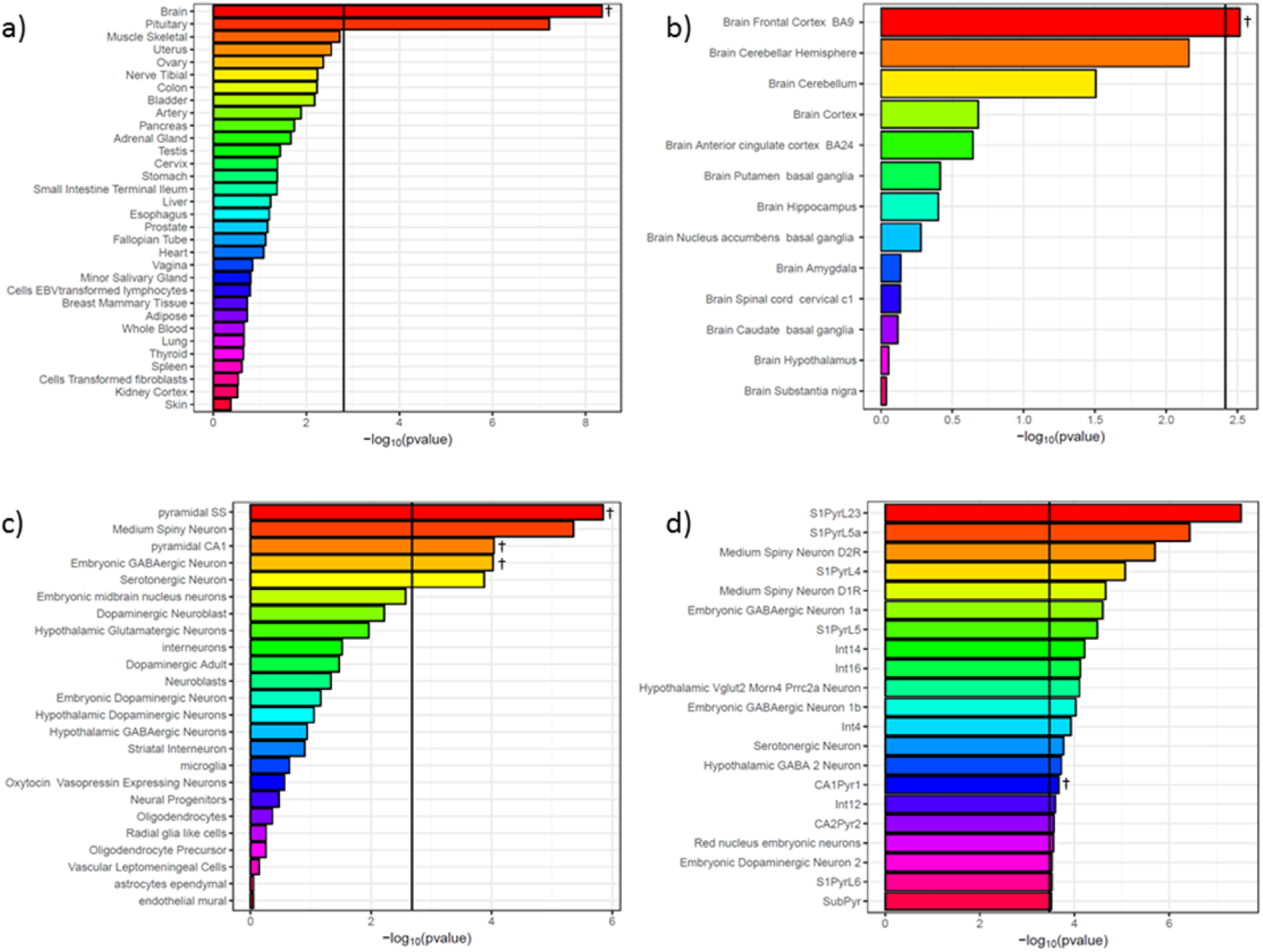
Results of tissue- and cell-specific analyses. a) Whole-body analyses; b) Brain tissue analyses; c) KI level 1 cell analyses; d) KI level 2 cell analyses (only those significant in MAGMA competitive analyses after correction for multiple testing shown). Vertical lines are the Bonferroni threshold for each analysis. Results shown are from MAGMA analyses - tissues and cell-types also significant in LDSC analyses are indicated with †. Full results are shown in Supplementary Table 3.

In the KI level 1 (broad cell groups) cell-type specific analyses, both linear regression (MAGMA)^22^ and heritability enrichment-based analyses (LD Score)^23^ supported enrichment of genes with high specificity to pyramidal neurons in the somatosensory neocortex (p_MAGMA_ = 1.41 × 10^-6^, p_LDSC_ = 5.81 × 10^-4^) and in the CA1 region of the hippocampus (p_MAGMA_ = 9.08 × 10^-5^, p_LDSC_ = 1.12 × 10^-3^), as well as to midbrain embryonic GABAergic neurons (p_MAGMA_ = 9.47 × 10^-5^, p_LDSC_ = 1.61 × 10^-3^; Figure 3c, Supplementary Table 3c). Level 2 analyses (narrowly-defined cell types) suggested significant enrichment of genes with high specificity to type pyramidal cells of the CA1 region (p_MAGMA_ = 2.16 × 10^-4^, p_LDSC_ = 1.19 × 10^-4^), although considerable variability was observed between methods at this level of granularity (Figure 3d, Supplementary Table 3d).

Cell-type specific analyses were repeated conditioning on each enriched cell-type in turn. When controlling for gene expression in pyramidal neurons of the somatosensory neocortex, the previously observed enrichment in CA1 pyramidal cells is lost. In contrast, the patterns of enrichment observed when conditioning on expression in CA1 pyramidal cells or in midbrain embryonic GABAergic neurons were consistent with those observed without conditioning (Figure 4).

**Figure 4:**
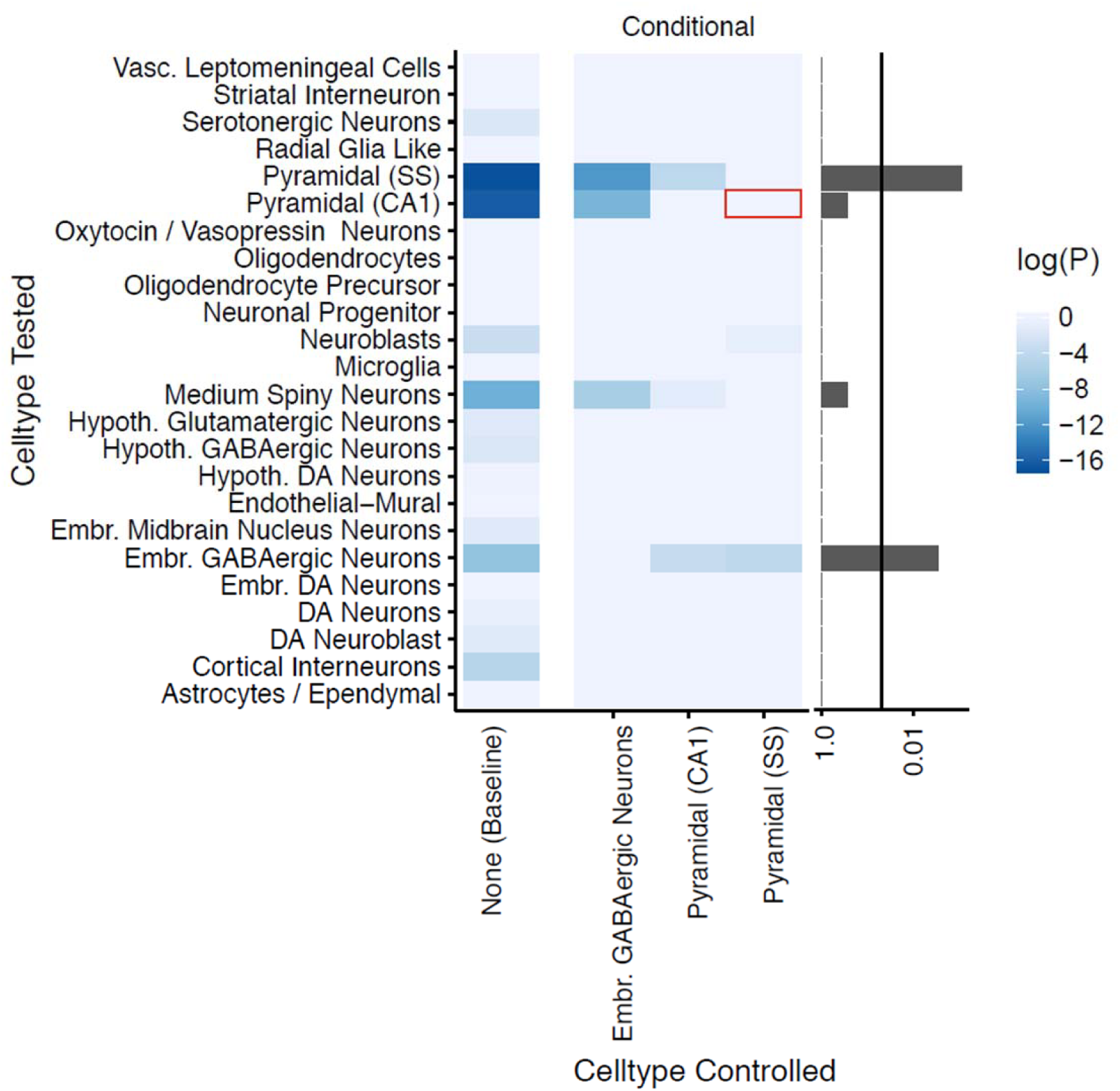
Conditional cell-type enrichment analysis. X-axis lists target cell-types. Y-axis lists other cell-types. Colors correspond to the enrichment probability of the other cell type after conditioning (P). Values of log(P) approaching zero indicate no enrichment after conditioning. Barplot on the right shows the minimum value of P for each cell-type across all conditional analyses (excluding analyses of the target cell-type with itself); the vertical line marks p=0.05. Red box highlights the loss of significant enrichment in CA1 pyramidal neurons when conditioning on somatosensory (SS) neurons.

### Predicted tissue-specific gene expression

Across the 10 GTEx brain tissues, results from MetaXcan suggested significant effects on the expression of 16 genes (p < 1.60×10^-6^; Table 3). Eight of these genes are situated at locus 11 (Chr3, 48.7-50.2 Mb). Genetic variation was predicted to upregulate 6 genes and downregulate 10 genes, with both single-region (9 genes) and multiple-region (7 genes) patterns of altered expression implied. Three genes (*NAGA, TUFM* and *GMPPB*) have previously been implicated in ID.

**Table 3:** Genes predicted to be upregulated (+) or downregulated (-) in specific tissues. Genetic locus is that identified in single-variant analyses, or the genomic region otherwise. Genewise = genome-wide significant in gene-wise analyses. Genes in italics have previously been implicated in developmental delay or intellectual disability.

| Locus | Gene | Tissues | Direction | P | Genewise? |
| --- | --- | --- | --- | --- | --- |
| 11 | RNF123 | Putamen basal ganglia<br>Nucleus accumbens basal ganglia<br>Frontal cortex<br>Cerebellar Hemisphere | -<br>-<br>-<br>- | $5.77 \times 10^{-7}$<br>$3.98 \times 10^{-10}$<br>$1.84 \times 10^{-7}$<br>$6.37 \times 10^{-7}$ | No |
| 11 | NPRL2 | Cortex | + | $1.06 \times 10^{-7}$ | No |
| 11 | MST1 | Hypothalamus | + | $1.33 \times 10^{-7}$ | Yes |
| 11 | MST1R | Cerebellum<br>Cerebellar Hemisphere | -<br>- | $2.25 \times 10^{-7}$<br>$8.98 \times 10^{-7}$ | Yes |
| 11 | RBM6 | Frontal cortex | + | $2.28 \times 10^{-7}$ | Yes |
| 11 | FAM212A | Cerebellar Hemisphere | - | $4.40 \times 10^{-7}$ | Yes |
| 11 | AMIGO3 | Cortex | - | $1.31 \times 10^{-6}$ | Yes |
| 11 | *GMPPB* | Cortex | - | $1.38 \times 10^{-6}$ | Yes |
| Chr12: 58.3-58.4 Mb | XRCC6BP1 | Nucleus accumbens basal ganglia<br>Hypothalamus<br>Cortex<br>Caudate basal ganglia | +<br>+<br>+<br>+ | $1.15 \times 10^{-6}$<br>$1.50 \times 10^{-6}$<br>$9.27 \times 10^{-7}$<br>$9.33 \times 10^{-7}$ | No |
| 15 | FOXO3 | Nucleus accumbens basal ganglia | - | $4.93 \times 10^{-10}$ | Yes |
| 23 | *TUFM* | Putamen basal ganglia<br>Nucleus accumbens basal ganglia | -<br>- | $4.89 \times 10^{-7}$<br>$1.38 \times 10^{-7}$ | Yes |
| 23 | NPIPB6 | Caudate basal ganglia | - | $1.33 \times 10^{-6}$ | No |
| Chr17:34.9-35.0 Mb | DHRS11 | Putamen basal ganglia<br>Nucleus accumbens basal ganglia<br>Caudate basal ganglia | -<br>-<br>- | $3.75 \times 10^{-7}$<br>$1.39 \times 10^{-6}$<br>$1.09 \times 10^{-6}$ | No |
| Chr18:12.9-13.2Mb | CEP192 | Cortex | + | $9.41 \times 10^{-8}$ | No |
| Chr22:42.4-42.5 Mb | *NAGA* | Putamen basal ganglia<br>Cerebellum<br>Cerebellar Hemisphere | -<br>-<br>- | $1.29 \times 10^{-7}$<br>$2.67 \times 10^{-7}$<br>$2.19 \times 10^{-7}$ | No |
| Chr22:42.5-42.6 Mb | CYP2D6 | Hypothalamus | + | $5.70 \times 10^{-7}$ | No |

### Tissue-specific pathway analysis

Tissue-specific pathway analyses identified 32 nested pathways with p ≤ 5.34×10^-6^ (Bonferroni correction for 9,361 effectively independent pathways), of which 29 contained genes that were significant in the gene-wise analysis (Supplementary Table 4; Supplementary Figure 2). 7 pathways remained significant after further correcting for all tissue-specific tests: “self-reported educational attainment”, “modulation of synaptic transmission”, “neurodegenerative disease”, “neuron spine”, “schizophrenia”, “rare genetic neurological disorder”, and “potentially synaptic genes” (Table 4).

**Table 4:** Pathways significantly enriched for genes associated with intelligence after correction for all tissue-effective pathways tests. Genes in italics have previously been implicated in developmental delay or intellectual disability.

| Pathway Name | Tissues significant | Significant genes | Min p [Tissue] |
| --- | --- | --- | --- |
| Self Reported Educational Attainment | Adipose, Artery, Bladder, Brain, Breast Mammary Tissue, Cells Transformed fibroblasts, Cervix, Colon, Fallopian Tube, Liver, Lung, Nerve Tibial, Ovary, Unweighted, Small Intestine Terminal Ileum, Spleen, Uterus, Whole Blood, Amygdala, Anterior cingulate cortex (BA24), Caudate (basal ganglia), Cortex, Frontal Cortex (BA9), Hippocampus, Hypothalamus, Nucleus accumbens (basal ganglia), Putamen (basal ganglia), Spinal cord (cervical c1) | ATXN2L, RNF123, NEGR1, DPP4, BTN1A1, JMJD1C | $1.03 \times 10^{-9}$<br>[Frontal Cortex (BA9)] |
| GO:Neuron Spine | Artery, Bladder, Cells EBVtransformed lymphocytes, Cervix, Esophagus, Lung, Minor Salivary Gland, Unweighted, Uterus, Vagina, Cerebellar Hemisphere, Cerebellum, Hypothalamus | ARFGEF2, SHANK3, EEA1, GRM3 | $5.29 \times 10^{-8}$<br>[Cerebellum] |
| GO:Modulation Of Synaptic Transmission | Artery, Bladder, Cells EBVtransformed lymphocytes, Cervix, Esophagus, Ovary, Thyroid, Uterus, Vagina, Hippocampus | STAU1, PLCL2, DBN1, SHANK3, SHISA9, GRM3, IGSF9B | $2.77 \times 10^{-8}$<br>[Uterus] |
| Schizophrenia | Brain, Pituitary, Testis, Cortex, Frontal Cortex .BA9. | FOXO3, ARFGEF2, PLCL2, DBN1, APBA1, AMT, THRB, GRK6, FOXO6, SHANK3, GPX1, DCC, EXOC4, F12, CDC42, SHISA9, NEGR1, GRM3, DPP4, IGSF9B | $5.57 \times 10^{-8}$<br>[Pituitary] |
| Rare Genetic Neurological Disorder | Brain, Nerve Tibial, Pituitary | FOXO3, CSE1L, ARFGEF2, PDE1C, DBN1, AMT, THRB, GRK6, ZNF638, FOXO6, RHOA, SHANK3, ATXN2L, DAG1, TRAIP, BSN, EEA1, GPX1, DCC, TUFM, DHODH, EXOC4, F12, GMPPB, MARCH5, SH2B1, MST1R, DYSF, TSHZ3, COL11A1, PRKAG1, KMT2D, CDC42, ATP2A1, DDN, WNT4, ACR, APEH, GRM3, DPP4, CYSTM1, SEMA3F, JMJD1C, TET2, CKB | $9.09 \times 10^{-8}$<br>[Brain] |
| Neurodegenerative Disease | Testis | FOXO3, CSE1L, *ARFGEF2*, PDE1C, DBN1, APBA1, *THRB*, GRK6, ZNF638, *SHANK3*, ATXN2L, *DAG1*, BSN, EEA1, GPX1, DCC, *TUFM*, EXOC4, F12, MARCH5, DYSF, *COL11A1*, *KMT2D*, CDC42, ATP2A1, ACR, APEH, NEGR1, GRM3, DPP4, CYSTM1, SEMA3F, JMJD1C, VWA5A, SEH1L | $4.75 \times 10^{-8}$<br>[Testis] |
| Potentially Synaptic All | Brain | FOXO3, CSE1L, *ARFGEF2*, PDE1C, PLCL2, DBN1, APBA1, *THRB*, ZNF638, FOXO6, RHOA, *SHANK3*, ATXN2L, EFTUD1, BSN, EEA1, GPX1, DCC, *TUFM*, PREX1, EXOC4, IP6K1, SH2B1, CAMKV, PRR7, ARHGAP15, TSHZ3, RBM6, *KMT2D*, CDC42, SHISA9, ATP2A1, DDN, SEPT3, NEGR1, GRM3, USP4, DPP4, DCAF5, IGSF9B, JMJD1C, TET2, VWA5A, SEH1L, PITPNM2, CKB | $1.16 \times 10^{-7}$<br>[Brain] |

### Conditional gene set enrichment

Results from tissue-specific and gene-set analyses identified a number of gene sets associated with intelligence. Of specific interest were synaptic genes (post-synaptic density proteome list;^24^), RBFOX family binding partners^25^, CELF4 binding partners, and previously reported intellectual disability genes. Additional analyses were performed to assess whether the association of these gene sets was independent of the enrichment for gene expression in pyramidal cells of the somatosensory cortex. Bootstrapped analyses confirmed the association of each gene set prior to conditional analysis (empirical p < 0.05; Supplementary Table 5). Conditioning on expression in gene expression in pyramidal cells of the somatosensory cortex, the enrichment for synaptic genes is no longer significant, indicating that this enrichment is not independent of that observed in the pyramidal neurons. This effect was not observed for RBFOX or CELF4 targets. Intellectual disability genes were weakly enriched prior to conditioning, and conditioning had little effect on enrichment. However, subdividing ID/DD genes into those associated with severe and those associated with moderate ID/DD indicated that the enrichment stemmed predominantly from moderate ID/DD genes, and this was not altered by conditioning on somatosensory pyramidal neuron gene expression (Supplementary Table 5).

## Discussion

The overall power of our GWAS meta-analysis was equivalent to a sample size of ∼99,000 individuals due to the inclusion of the extreme trait HiQ sample, which contributes equivalently to a population cohort of ∼21,000 individuals, and is likely to be enriched for alleles associated with intelligence in the normal range. We mapped results from our GWAS to tissue and cell-type specific gene expression data, identifying enrichment of specificity at multiple levels: in the brain, the frontal cortex, midbrain embryonic GABAergic neurons and pyramidal neurons, especially those in the somatosensory cortex. A number of genes previously implicated in intellectual disability or developmental delay are predicted from the GWAS results to show differential gene expression for normal range IQ in different brain regions, and are associated with variation in intelligence in the normal range from gene-wise analyses.

RNA sequencing data suggest that genes more strongly associated with intelligence are enriched for brain-specific expression in general. While the dominance of brain-specificity over other body tissue specificity is pronounced, assessing differences within the brain is more difficult. Genes more strongly associated with intelligence showed higher specificity for frontal cortical expression, but the differences in cell composition between brain tissues means that cell-type analyses may be more informative. This can be seen within our results, in that pyramidal neurons of the somatosensory cortex were significant in the cell-type specific analysis, but the cortex as a whole is not significant in the GTEx brain-tissue analysis. This is perhaps due to the fact that the cortex is a highly heterogeneous mixture of cell-types. Our results suggest expression in pyramidal neurons in one area of the cortex is relevant in intelligence, but expression in the other cell-types and areas of the cortex may not be^26^. A further caveat to this interpretation is that the full cellular composition of the cortex (and the brain overall) is not reflected in the KI mouse superset, and as such our conclusions about wider effects must be constrained.

Genes with higher specificity to pyramidal neurons were enriched for associations with intelligence. However, the location of the most interesting neuron population is not yet clear. Observed enrichment in the pyramidal cells of the CA1 region of the hippocampus was lost when accounting for gene expression in pyramidal cells of the somatosensory cortex. An uncaptured population of pyramidal neurons (for example, in the frontal cortex) may exist that similarly overlaps in expression with the somatosensory pyramidal neurons, and accounts for the enrichment observed in the latter population. While the KI superset is the largest brain scRNAseq resource to date, it covers a limited set of regions and developmental stages^16^.

Tissue and cell-specific analyses were performed using related but distinct approaches, namely linear regression of binned specificity and heritability enrichment analysis of the top 10% of specific genes. Linear regression tended to give smaller p-values than heritability enrichment analysis (Supplementary Table 3). While this may be a statistical artefact generated by the differing assumptions of the methods, this could also result from the pattern of enrichment. For example, a small set of genes associated with intelligence, all with very high specificity to a given tissue would result in a lower p-value from heritability enrichment than if the association with intelligence was spread more broadly across genes with generally enriched tissue-specificity.

The results of analyses in the KI mouse superset for intelligence can be contrasted with those for schizophrenia^16^. Both phenotypes initially demonstrate enrichment for the same hippocampal and somatosensory pyramidal neuron populations. However, conditional analyses in schizophrenia demonstrated significant differences in the results between schizophrenia and intelligence - the enrichment observed in the somatosensory cortex in schizophrenia could be fully explained by that in the hippocampus, while the opposite pattern was observed in intelligence. Additionally, unlike in schizophrenia, genes implicated in intelligence also showed specificity in midbrain embryonic cells. Analyses in schizophrenia implicated more level two cell types than were significant in our analyses. Such differences may reflect differences in the common-variant contribution to variance in intelligence and schizophrenia; despite a similar effective sample size to the schizophrenia GWAS (40,675 cases, 64,643 controls, N_eff_ ∼ 100K), this GWAS identified only 25 loci compared with the 140 associated with schizophrenia^27^ highlights the potentially more polygenic nature of the observed heritability of intelligence, which is approximately equal to that of schizophrenia.

The KI mouse superset has a number of strengths; it is the largest and broadest scRNAseq dataset to date, captures extra-nuclear as well as nuclear transcripts, and was generated using identical methods^16^. However, there are also limitations to the use of this dataset. One issue is that the expression data used is derived from mice rather than humans. Gene expression in the brain is conserved across mammals, such that the principal axes of variation in comparative studies of gene expression capture inter-tissue, rather than inter-species, variance^28^. Furthermore, there is a high degree of conservation of gene expression between mouse and human brains specifically^16,29,30^. Previous analyses using the KI mouse superset have made extensive comparison between mouse and human gene expression and found high concordance in mapping mouse to human genes^16^.

Nevertheless, cell types that are not enriched for genes associated with intelligence in this study should not be prematurely excluded, as some cell-types are not present in the dataset and others will have dissimilar functions or have been exposed to different evolutionary pressures in mouse and in human. Intelligence is a major characteristic that differentiates humans from other mammals^31^. As such, it may be the case that genes with higher specificity to regions dissimilar between humans and mice could be enriched for associations with intelligence, which would not be captured by this approach.

Our results highlight potential insights beyond those gleaned from tissue-specific expression patterns. Several genes previously implicated in ID/DD were present in loci associated in the GWAS. Perhaps the most interesting example of this is *GMPPB* (GDP-Mannose Pyrophosphorylase B), which is in locus 11 of the GWAS results, and was genome-wide significant in gene-wise analyses. It is a member of the “rare genetic neurological disorder” gene set (significantly enriched, specifically in neural tissues), and the expression of *GMPPB* was predicted to be downregulated in the cortex of individuals with higher intelligence. Rare loss of function mutations in the *GMPPB* gene have been identified as causal mutations in tens of individuals with muscular dystrophies and myasthenias, many of which present with mild to severe ID^32–36^. The product of this gene is important for the glycosylation of a-DG (alpha-dystroglycan - the dystroglycan gene *DAG* is also present in locus 11 and is a significantly associated gene in this analysis;^32^). Glycosylation is required for the interaction of a-DG with extracellular ligands, with a variety of consequences including the organization of axon guidance^37^. Our results, in the context of the medical genetic literature, tentatively suggest the effects on axon guidance by glycosylated a-DG may be an area worthy of further exploration to understand the biology of intelligence.

However, the observed overall overlap of all ID/DD genes with loci from the GWAS does not differ from that expected by chance. This lack of significant enrichment is partly reflected in the pathway analysis - there are several overlapping gene sets designed to capture ID/DD genes. Of these, only “rare genetic neurological disorder” was significantly enriched following correction. Further insight is obtained from conditional gene-set analyses - although the overall “intellectual disability” gene set is only nominally associated with intelligence, stratifying this gene set demonstrates considerable enrichment of mild intellectual disability genes, independent of gene expression in somatosensory pyramidal neurons. The presence of genes causative of ID/DD in loci associated with intelligence in the normal range, and the enrichment of specific pathways and gene sets derived from the ID/DD literature, may indicate shared biology between ID/DD and normal intelligence. However, the lack of broad enrichment of all such genes suggests that there may also be distinct pathways contributing to normal intelligence that are not commonly affected in ID/DD. ID/DD is not a single condition, but a group of disorders with differing etiologies - our results are still consistent with a two-group model of ID/DD etiology^38–40^.

The meta-analysis results presented herein extend previous findings^4^. The results are largely consistent with those previously reported, which is unsurprising. More genes were identified in our gene-wise analysis due to an analytical decision to extend the boundaries by which each gene is defined 35kb upstream and 10kb downstream of the coding region^41,42^. Defining genes using this boundary (as opposed to no boundary extension) captures additional transcriptional elements - these may be specific to the target gene, but may also capture elements with more distal regulatory effects.

Deriving testable biological hypotheses from the statistical associations of GWAS results is one of the central challenges for the immediate future of complex genetics^3,15^. The provision of high-quality reference datasets encompassing genetic information from variation to translation, and the integration of genomic data to such reference data is invaluable to this aim^16,18^. We have demonstrated that some insights into the biology of intelligence can be derived from GWAS, and have suggested potential avenues for further exploration. Our results could indicate that intelligence represents optimal pyramidal neuron functioning. Cognitive tests are highly correlated with general intelligence (*g*), which may depend on pyramidal neuron function^7^.

Understanding how biology underlies variation in intelligence is an active area of research that is beginning to yield results. Unifying these new genetic results with data from multiple approaches, can increase the power of each approach, has the potential to yield greater understanding of the biology of intelligence, which in turn could inform the study of many health-related phenotypes with which intelligence is correlated.

## Online Methods

### Cohort descriptions

#### Sniekers intelligence GWAS^4^

The cohort analyzed in Sniekers et al. (2017) was drawn from 7 cohorts, primarily consisting of data from the UK Biobank pilot genotyping release (N = 54,119) and the Childhood Intelligence Consortium (N = 12,441) as well as seven additional cohorts (N = 11,748). The phenotype for analysis was Spearman’s *g*, or a primary measure of fluid intelligence that correlates highly with *g*^4,43^. Summary statistics from this analysis are available at https://ctg.cncr.nl/software/summary_statistics. Full details on cohort characteristics, genotyping and analysis are supplied in the Supplementary Material.

#### HiQ high-intelligence GWAS

The Duke University Talent Identification Program (TIP) cohort has been described previously^17^. In brief, TIP is a non-profit organization that recruits and nurtures academically gifted children of extremely high intelligence (top 3%) from the US population. For genomic study, 1,247 participants from the top 1% of TIP (top 0.03% of population) were selected as a high-intelligence cohort (HiQ). IQ was inferred from performance on the Scholastic Assessment Test (SAT) or American College Test (ACT) taken at age 12 rather than the usual age of 18 years. A population comparison cohort (N = 8,185) was obtained from the The University of Michigan Health and Retirement Study (HRS). Participants were assumed to be drawn from the normal distribution of intelligence. Full details on genotyping and analysis are supplied in the Supplementary Material.

### Meta-analysis

Summary statistics from Sniekers et al. (2017) and HiQ were meta-analyzed using METAL^4,44^. To account for the increased discovery power afforded by the extreme-sampling method used in TIP, analyses were weighted by their respective non-centrality parameters (NCP), estimated using the Genetic Power Calculator^45–47^. Specifically, NCPs were estimated assuming a causal variant of 20% frequency, capturing 0.1% of variance in each phenotype, assuming HiQ controls are drawn from the normal distribution (+/- 2 SD from the mean) and HiQ cases are sampled from 4 SD above the mean, consistent with IQ > 160 in 99% of the cohort^17^. The NCP of the Sniekers cohort (N = 78,308) was 78.4, while the NCP of the HiQ cohort was 21.6, suggesting the HiQ cohort contributes equivalently to a population cohort of ∼21,000 individuals. Only variants present in both cohorts were retained for analysis.

Following association analysis, genome-wide significant loci were defined via clumping in PLINK2^48^. Index variants (p < 5×10^-4^) were merged into loci if in linkage disequilibrium (r^2^ > 0.1 within 500kb) with a variant with a lower p-value. Loci within 50kb of each other were merged. Manhattan and QQ plots were generated using FUMA (^49^. Annotation of genomic results with: data from the EBI GWAS catalog; OMIM; GENCODE genes; genes previously implicated in autism and in intellectual disability; copy-number variants previously implicated in psychiatric disorders; and mouse knockout phenotypes was performed with RegionAnnotator version 1.63 (https://github.com/ivankosmos/RegionAnnotator).

### Heritability and partitioned heritability

The heritability of intelligence accounted for by common variants was estimated using LD Score, limited to the HapMap3 variants and pre-computed LD scores provided with the package^19^. Heritability was then partitioned across the 53 genomic annotations provided with the package^23^, with the addition of 5 annotations: open chromatin regions (ATAC and ATAC Bryois extend 500, which increases the window around the region by 500 bases in both directions), the intersection between ATAC and conserved regions of the genome (ATAC-conserved) and regions present in the Neanderthal genome (Neanderthal and Neanderthal extend 500;^21,50^. Regions of open chromatin were identified in prefrontal cortical tissue from 135 schizophrenic individuals and 137 controls using ATAC sequencing, which identifies stretches of DNA free of nucleosomes and other DNA-binding proteins^21,51^.

### Gene-wise analyses

Results from the meta-analysis were filtered to retain only single nucleotide variants (SNPs) present in the European superpopulation of 1000 Genomes Phase 3^52^. SNPs were annotated to a gene using MAGMA v1.06, assigning SNPs to genes if they lay between 35kb upstream and 10kb downstream of the gene location (as supplied on the MAGMA website, build 37;^22^. Gene-wise p-values were obtained from MAGMA as the aggregate of the mean and smallest p-value across all SNPs annotated to the gene. MAGMA accounts for possible confounders such as gene size, gene density, linkage disequilibrium and minor allele count. The threshold for genome-wide significance was defined as p = 2.65 × 10^-6^, the Bonferroni correction for the 18,839 genes tested. Genes passing genome-wide significance were defined as coming from the same locus if their locations were within 50kb of each other, or if they lay within clumped loci from the single variant analysis. Significant genes were cross-referenced to the intellectual disability (ID) gene list provided with RegionAnnotator. The significance of the observed overlap was quantified as a hypergeometric test in R^53^, using as background 1,366 ID/DD genes in 18,839 autosomal genes.

### Tissue- and cell-specific gene expression

Tissue-specific and cell-type specific proportions of gene expression were calculated following the method described in detail in^16^). Tissue expression data was drawn from the GTEx Consortium^18^, and brain cell-type expression data was drawn from scRNAseq data from mouse brain^16^. For each gene, the value for each tissue (or cell-type) was calculated by dividing the mean Unique Molecular Identifier (UMI) counts for the given tissue by the summed mean UMI counts across all tissues^16^.

Associations between gene-wise p-values from the meta-analysis and tissue-specific (cell-type specific) gene expression were calculated using two methods, implemented in MAGMA^22^ and in LD Score^19^. In MAGMA, genes were grouped into 40 equal bins by specificity of expression, and bin membership was regressed on gene-wise association with intelligence in the meta-analysis (For these analyses, gene-wise association was defined as the mean p-value across all SNPs assigned to the gene.) In LD Score, the 10% of genes with the highest specificity within each tissue were used as a gene set for partitioned heritability analysis. Results were considered significant if the association p-values were smaller than the relevant Bonferroni threshold for both methods.

Conditional cell-specific analyses were performed as a secondary analysis to test whether each enriched cell-type observed was independent of all others. Full details of the method implemented are provided in Skene et al., 2017. In brief, for each enriched cell-type in turn (the target cell-type), z-scores from gene-wise association analyses with intelligence were randomly resampled without replacement. The mean z-score within each expression-specificity decile of the target cell-type was held constant, but the mean z-score of each specificity decile of other cell types was randomized. Empirical p-values were derived for each of the other cell-types, and this procedure was repeated 10,000 times. Expression in each of the other cell-types is considered to be associated with intelligence independently of expression in the target cell type if the observed p-value is lower than the 500th empirical p-value (i.e. 95% of the empirical distribution;^16^).

### Predicted tissue-specific gene expression

Results from the variant-level meta-analysis were used to predict gene expression using MetaXcan and genomic and transcriptomic reference data from the brain regions assayed in the GTEx project^18,54^. Associations between predicted gene expression levels and intelligence were calculated. Significance was set at 1.60×10^-6^, the Bonferroni correction for the 31303 gene-tissue pairs tested^54^. Significant genes were cross-referenced to the intellectual disability gene list provided with RegionAnnotator.

### Pathway analysis

A pathway matrix **P** was generated with elements *P_g,p_* = 1 if gene *g* was in pathway *p* and *P_g,p_* = 0 otherwise. The elements in the matrix were multiplied by binned gene expression weights obtained from GTEx data (as in the foregoing section on tissue-specific expression) for 13 brain regions and 32 tissues, generating 45 weighted gene/pathway matrices. Only genes with expression data were taken into account. Association between these tissue-weighted pathways and gene-wise associations with intelligence was computed using MAGMA. 13,564 pathways were drawn from OpenTargets (downloaded January 2017;^55^), GO ontologies, canonical pathways drawn from MSigSB v5.2 C2 and C5 datasets^56^, and biological pathways related to psychiatric disorders found in various scientific publications (a link to each pathway source is provided in Supplementary Table 4). The pathways assessed by this approach are related to each other in a complex fashion; GO pathways are hierarchical and MSigDB and OpenTargets pathways capture related gene sets. Accordingly, in order to control for multiple testing, the effective number of pathways tested was established by computing the number of principal components accounting for 99.5% of explained variance in the pathway similarity matrix, obtained by computing the Tanimoto similarity between pathways. This results in a Bonferroni-corrected threshold of p = 5.34×10^-6^ for 9,361 effectively independent tests for each matrix. A more stringent threshold was applied secondarily, taking into account all tissue-specific pathway matrices for a total for 9,361 × 46 tests and threshold p = 1.16×10^-7^.

### Conditional gene set enrichment

Gene sets of interest were drawn from the results of pathway analyses. Specifically, the human postsynaptic density proteome gene set^24^ was used to capture the effects of synaptic and neuronal pathways. Gene sets of RBFOX and CELF4 targets were included as they showed enrichment that appears to be driven by brain-specificity, and have been previously implicated in other brain-related traits^16,25,57^. ID/DD gene sets were tested as they are of specific interest to the study of intelligence. To assess whether the enrichment of these gene sets is independent of gene expression in somatosensory pyramidal neurons, conditional analyses were performed following the method described above for conditional cell-type analyses, modified for the use of gene sets^16^.

## Data availability

Summary statistics from the GWAS meta-analysis will be made available at [link available upon publication].

## Acknowledgements

We gratefully acknowledge the contribution of all of the researchers and participants involved in the collection and analysis of the data included. This study includes data from Sniekers et al., 2017, which made use of the UK Biobank resource under application number 16406 (as previously acknowledged).

Analysis in this paper represents independent research funded by the National Institute for Health Research (NIHR) Biomedical Research Centre at South London and Maudsley NHS Foundation Trust and King’s College London. The views expressed are those of the authors and not necessarily those of the NHS, the NIHR or the Department of Health. Analyses were performed using high performance computing facilities funded with capital equipment grants from the GSTT Charity (TR130505) and Maudsley Charity (980).

Analysis from Sniekers et al., 2017 was funded by the Netherlands Organization for Scientific Research (NWO VICI 453-14-005), and carried out on the Genetic Cluster Computer, which is financed by the Netherlands Scientific Organization (NWO:480-05-003), by VU University, Amsterdam, the Netherlands, and by the Dutch Brain Foundation and is hosted by the Dutch National Computing and Networking Services SurfSARA. PRJ is supported by the ‘Stichting Vrienden van Sophia’ (grant nr: 14-27) awarded to DP.

Research on the HiQ cohort was supported by a European Research Council Advanced Investigator award (295366) to RP. Collecting DNA from the highest-scoring TIP individuals was supported by an award from the John Templeton Foundation (13575) to RP.

Summary statistics from this analysis have been made available at [link available upon publication].

## Author contributions

GB, DP and PFS conceived the study. JRIC, JB, HAG, PRJ, JS and NS performed statistical analyses. RP, ABM, SL, GC and JH acquired data. JRIC and GB wrote the manuscript. All authors reviewed the manuscript.

